# Molecular basis for a germline-biased neutralizing antibody response to SARS-CoV-2

**DOI:** 10.1101/2020.11.13.381533

**Authors:** Sarah A. Clark, Lars E. Clark, Junhua Pan, Adrian Coscia, Lindsay G.A. McKay, Sundaresh Shankar, Rebecca I. Johnson, Anthony Griffiths, Jonathan Abraham

## Abstract

The SARS-CoV-2 viral spike (S) protein mediates attachment and entry into host cells and is a major target of vaccine and drug design. Potent SARS-CoV-2 neutralizing antibodies derived from closely related antibody heavy chain genes (IGHV3-53 or 3-66) have been isolated from multiple COVID-19 convalescent individuals. These usually contain minimal somatic mutations and bind the S receptor-binding domain (RBD) to interfere with attachment to the cellular receptor angiotensin-converting enzyme 2 (ACE2). We used antigen-specific single B cell sorting to isolate S-reactive monoclonal antibodies from the blood of a COVID-19 convalescent individual. The seven most potent neutralizing antibodies were somatic variants of the same IGHV3-53-derived antibody and bind the RBD with varying affinity. We report X-ray crystal structures of four Fab variants bound to the RBD and use the structures to explain the basis for changes in RBD affinity. We show that a germline revertant antibody binds tightly to the SARS-CoV-2 RBD and neutralizes virus, and that gains in affinity for the RBD do not necessarily correlate with increased neutralization potency, suggesting that somatic mutation is not required to exert robust antiviral effect. Our studies clarify the molecular basis for a heavily germline-biased human antibody response to SARS-CoV-2.

## Introduction

The SARS-CoV-2 pandemic has caused over 48 million infections with more than 1.2 million deaths worldwide. Vaccines and therapeutic agents are urgently needed, particularly as certain parts of the world now face new waves of infections. The SARS-CoV-2 spike protein (S) is a large and heavily glycosylated protein that forms trimers of heterodimers on the surface of virions. Each S protomer has two functional subunits; S1, which contains a receptor-binding domain (RBD) that binds to ACE2^1,2^, and S2, which mediates fusion of the viral and host cell membranes during viral entry.

Epitopes for neutralizing antibodies on S include sites on the RBD and on the S1 N-terminal domain (NTD)^3–8^. IGHV3-53 or IGHV3-66 antibody genes are identical except for a single amino acid mutation in an antibody framework region (FWR)^9^, and potent SARS-CoV-2 neutralizing antibodies derived from these two germline genes have been isolated from multiple COVID-19 convalescent individuals^3,4,8,10–13^. The S RBDs can be in “down” or “up” conformations^14,15^, and ACE2 and IGHV3-53/3-66 neutralizing antibodies can only bind the RBD when it is “up” ^8,16^. Although described IGHV3-53/3-66 neutralizing antibodies have short CDR H3 loops, some IGHV3-53 antibodies with longer CDR H3 loops can make contacts with neighboring RBDs to close the trimer^17^.

Here, we used single B-cell sorting to isolate a panel of monoclonal antibodies that react against the SARS-CoV-2 S protein from the memory B cells of a COVID-19 convalescent individual. The most potent neutralizing antibodies were seven somatically related variants of a single IGVH3-53-derived antibody that binds the RBD with varying affinity. We use X-ray crystal structures of Fab/SARS-CoV-2 RBD complexes to explain the basis for gains or losses in RBD affinity that occurred during antibody maturation. We show that a germline revertant of the antibody binds tightly to the SARS-CoV-2 RBD and that gains affinity for the RBD are not associated with higher neutralization potency. We propose that such a focused germline-biased antibody to SARS-CoV-2 may be particularly vulnerable to antibody neutralization escape as the virus continues to circulate in humans.

## Results

### Related antibodies dominate the SARS-CoV-2 B-cell response in a convalescent donor

To study neutralizing antibody responses to SARS-CoV-2, we obtained a peripheral blood sample from a healthy individual (“C1”) who had been infected by SARS-CoV-2 five weeks prior to sampling. Polyclonal immunoglobulin G (IgG) purified from the blood of this individual neutralized SARS-CoV-2 but not vesicular stomatitis virus (VSV) lentivirus pseudotype (Fig. 1a). We generated a soluble SARS-CoV-2 S construct that is stabilized through mutations and the addition of trimerization tag to adopt the S “pre-fusion” conformation (“S2P”)^14^ and used it as an antigen to isolate 116 memory B cells (CD19^+^, IgG^+^) by FACS (Supplementary Fig. 1a). We could produce 48 recombinant monoclonal antibodies in sufficient amount for further characterization. Forty-three of these antibodies bound S2P by ELISA, and 18 also bound the RBD (Supplementary Fig. 1b and Supplementary Table 1). Most antibodies were derived from the IGHV3 heavy chain subgroup and had kappa light chains (Fig. 1b). Antibody CDR H3 and CDR L3 loops had an average length of 15 and 9 amino acids, respectively, with low frequencies of somatic hypermutation in variable heavy and light chain sequences (Fig. 1c-d and Supplementary Table 1).

**Figure 1.**
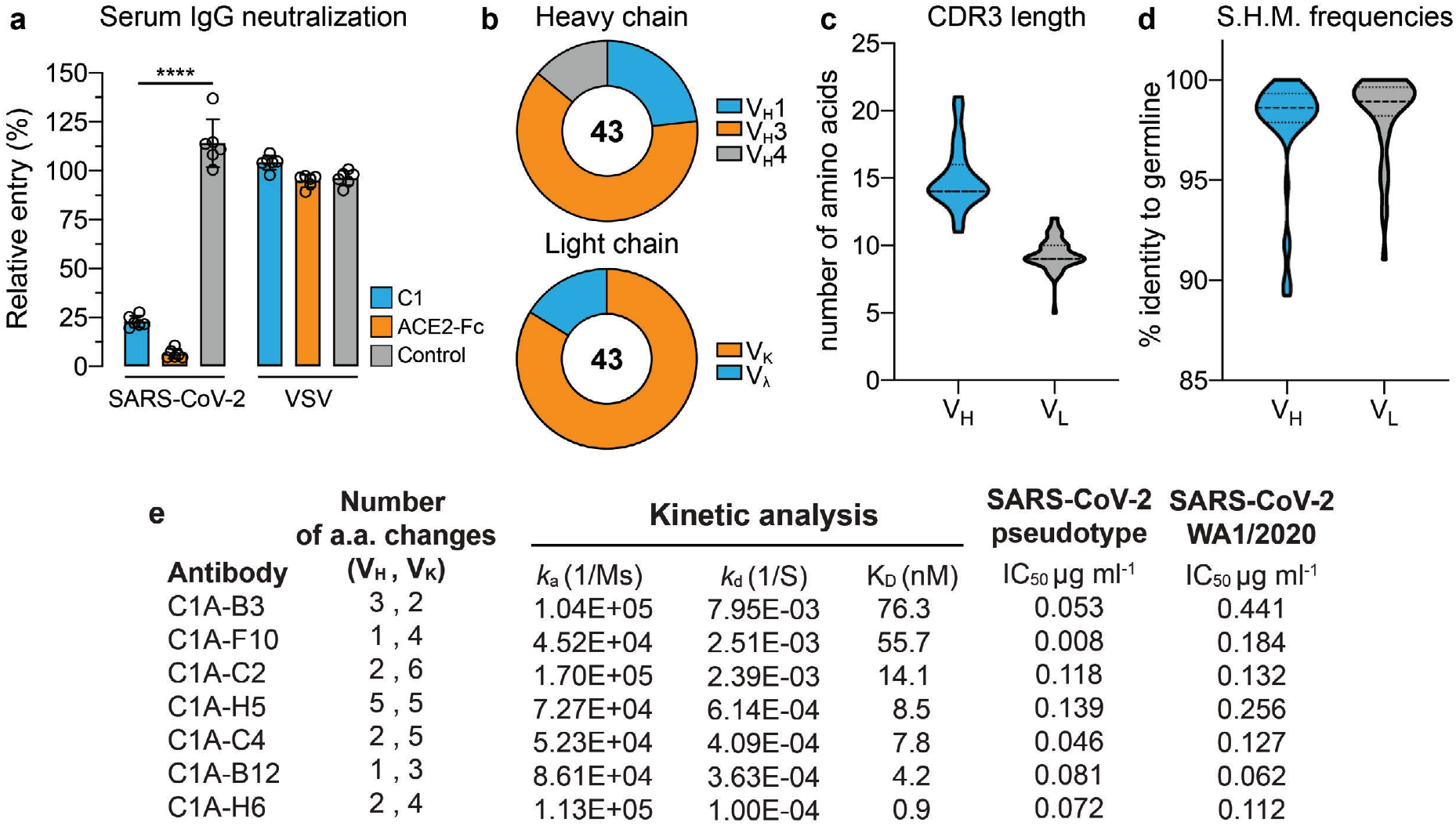
S-reactive monoclonal antibodies from a COVID-19 convalescent individual. (a) Entry levels of SARS-CoV-2 or vesicular stomatitis virus (VSV) pseudotypes after pre-incubation with polyclonal IgG purified from the plasma of a COVID-19 convalescent individual (“C1”), a non-immune control donor (“control”), or with an ACE2-Fc fusion protein. Data are normalized to a no antibody control. Means ± standard deviation from two experiments performed in triplicate (n=6). One-way ANOVA with Tukey’s multiple comparisons test. \*\*\*\**P* <0.0001. (b) Antibody heavy and light chain gene usage for SARS-CoV-2 S-reactive monoclonal antibodies. (c-d) Violin plots showing CDR3 loop lengths and somatic hypermutation frequencies (S.H.M.) for S-reactive monoclonal antibodies. The median and quartiles are shown as dashed and dotted lines, respectively. For CDR3 loop lengths, the median and first quartile marker overlap. (e) Properties of the seven IGHV3-53-derived potent SARS-CoV-2 neutralizing antibodies. a.a.: amino acids. WA1/2020: SARS-CoV-2 strain USA/WA1/2020.

Of the 43 antibodies we tested, only eight neutralized SARS-CoV-2 pseudotype with greater than 90% reduction in entry at a screening concentration of 100 μg ml^-1^ (Supplementary Fig. 1c). IC_50_ values ranged from 0.008 to 0.671 μg ml^-1^ in dose response pseudotype neutralization assays (Fig. 1e and Supplementary Fig. 2a). These eight antibodies also neutralized infectious SARS-CoV-2, but authentic virus was more resistant to antibody neutralization than pseudotype (Fig. 1e and Supplementary Fig. 2b).

The only antibodies that neutralized infectious SARS-CoV-2 with an IC_50_ value of less than 1 μg ml^-1^ – C1A-B3, -F10, -C2, -H5, -C4, -B12, and -H6 – were somatic variants of the same IGHV3-53/IGKV1-9-derived antibody (Supplementary Table 1). Each had a low number of amino acid substitutions in the heavy and light chain variable genes (Fig. 1e). Monomeric Fabs derived from these antibodies bound tightly to the RBD, with affinities ranging from 66 nM to 1 nM (Fig. 1e and Supplementary Fig. 3). C1A-B12, which we used as a representative member of the C1A-IGHV3 antibodies, prevented an ACE2-Fc fusion protein from binding to the RBD in a biolayer interferometry (BLI)-based competition assay (Fig. 2a). The Fab of CR3022, a human antibody that does not compete with ACE2-binding^18^, did not affect C1A-B12 Fab or ACE2-Fc binding to the RBD (Fig. 2a).

**Figure 2.**
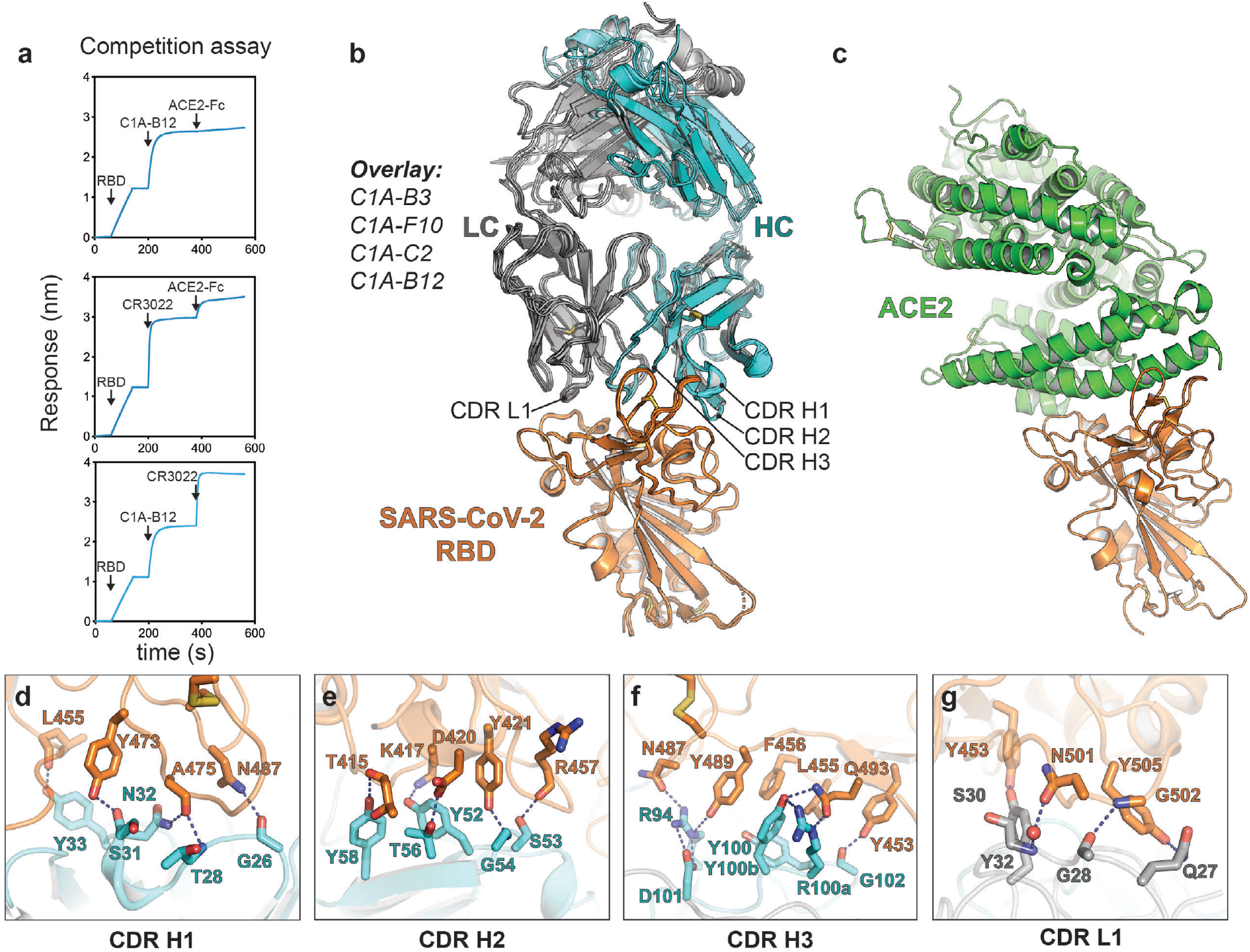
SARS-CoV-2 receptor-binding domain recognition by C1A-IGHV3-53 antibodies. (a) BLI-based competition assay for C1A-B12 Fab, CR3022 Fab, and human ACE2-ectdomain Fc fusion protein (ACE2-Fc) binding to the SARS-CoV-2 RBD. Arrows show the time point at which the indicated protein was added. Representative results of two replicates for each experiment are shown. (b) Overlay of ribbon diagrams for X-ray crystal structures of Fab/SARS-CoV-2 RBD complexes. CDR loops contacting the RBD are indicated. (c) Ribbon diagram of the X-ray crystal structure of the SARS-CoV-2 RBD bound to the ACE2 ectodomain (PDB ID: 6M0J)^44^ with the SARS-CoV-2 RBD in the same orientation as shown in *b* for comparison. (d-g) Details of the interface between the SARS-CoV-2 RBD and antibody CDR loops for antibody C1A-B3.

### Structures of Fab/SARS-CoV-2 RBD complexes

To better understand the effects of somatic mutations on RBD affinity, we determined X-ray crystal structures of the RBD bound to four Fabs: C1A-B3, -B12, -C2, and -F10 (Fig. 2b). The Fabs engage the RBD through an identical binding mode with root mean square deviations (r.m.s.d.) of 0.36-0.39 Å by structural superposition. As with other IGHV3-53/IGHV3-66-derived antibodies^4,8,10,12,19^, CDR loops H1, H2, H3, and L1 make the majority of RBD contacts (Fig. 2b). As suggested by the results of competition assays (Fig. 2a), the antibodies and ACE2 bind the same site on the RBD (Fig. 2c). Most of the contacts are polar and involve backbone and sidechain atoms on both sides of the interface (Fig. 2d-g). Somatic mutations in the C1A-IGHV3-3 antibodies occurred in CDR loops and FWRs, and in the structure, some (e.g., the F10S and S14F mutations in the light chain) are positioned far from the RBD and unlikely to impact antigen affinity (Fig. 3).

**Figure 3.**
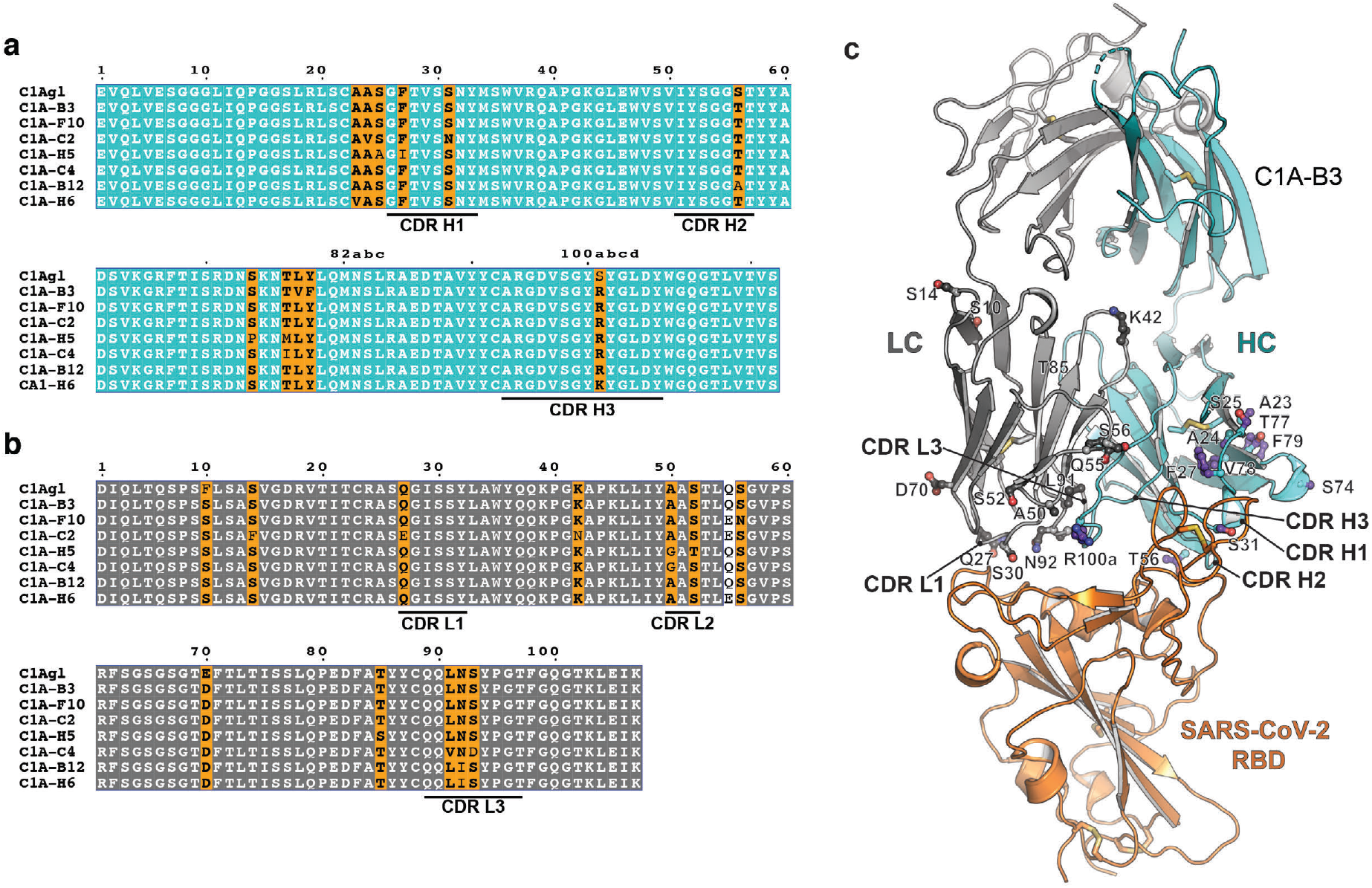
Positions of somatic changes on C1A-IGHV3-53-derived antibodies. Alignment of antibody variable heavy chain gene sequences (a) or variable light chain (b) gene sequences. C1A-gl are germline revertant sequences designed using IMGT/V-QUEST^31^. Panels were generated using ESPrit3^45^ and modified. The Kabat numbering scheme is used. (c) Ribbon diagram of crystal structure of the C1A-B3 Fab/RBD complex showing the location of somatic mutations.

### Structural basis for affinity maturation of IGHV3-53-derived antibodies

High-resolution X-ray crystal structures of multiple clonotypes allowed us to examine the effects of somatic mutations on the interaction interface. We also included in our analysis eight additional IGHV3-53/3-66-derived SARS-CoV-2 neutralizing antibodies isolated in other studies from multiple donors (B38, CC12.1, CC12.3, CV30, C105, BD-236, and BD-629)^4,8,10,12,19^. These antibodies have an essentially identical binding mode on the RBD (Supplementary Fig. S4).

As examples of mutations at the Fab/RBD interface mutations that likely increase affinity for the RBD, the V_H_ S31N and the S31R mutations, which are found in C1A-C2 and BD-629, respectively, provide new contacts with RBD Q474 and K458 (Fig. 4a-c). The V_H_ S56T mutation, which occurs in most of the C1A-IGHV3-53 antibodies (Fig. 3), positions a methyl group in van der Waals contact with RBD T415 and the side chains of Y52 and Y58 on the antibody (Fig. 4d-e). On the light chain, the N92I substitution creates a new hydrophobic contact with RBD Y505 (Fig. 4g-h). The V_H_ T28I somatic mutation, which we did not observe in C1A-IGVH3-53 antibodies, is probably important as it independently occurred in CV30^19^, B38^4^, and BD-629^8^ (Supplementary Fig. 5a). This change adds a hydrophobic contact with the Cα atom of RBD G476 and probably also helps orient CDR H1 to optimize neighboring polar contacts (Supplementary Fig. 5c-e). The residue at position 26 in the CDR H1 loop of IGHV3-53/3-66 antibodies is almost uniformly a glycine, but BD-629 contains a unique substitution (G26E) that provides a new set of polar contacts with the RBD (Supplementary Fig. 5e).

**Figure 4.**
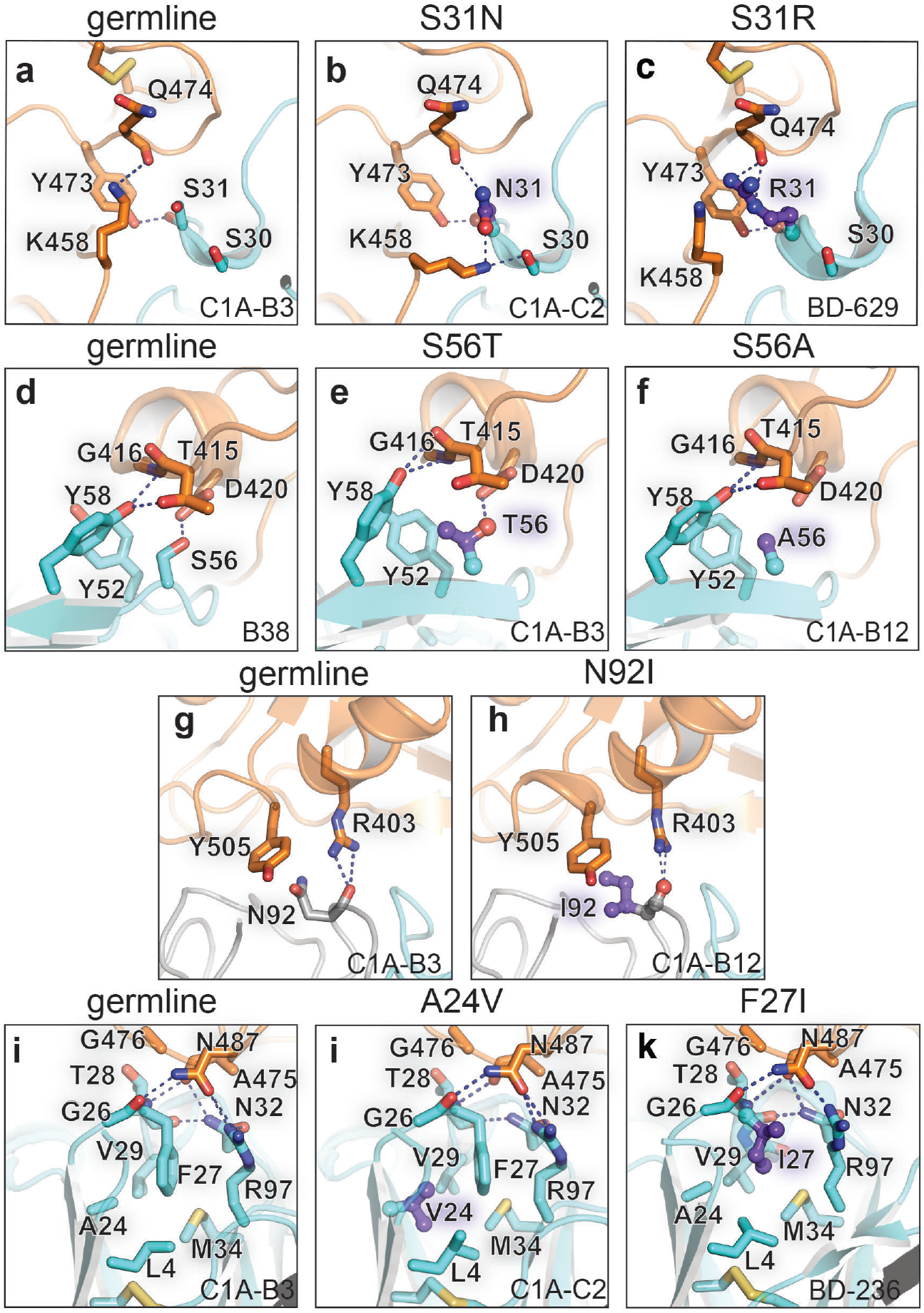
Antibody somatic mutations at the SARS-CoV-2 RBD interface. Interactions of CDR H1 residue 31 with the RBD are shown for C1A-B3 (a), C1A-C2 (b), or BD-629 (PDB: 7CH5)^8^ (c). Interactions of CDR H2 residue 56 with the RBD are shown for B38 (PDB: 7BZ5)^4^ (d), C1A-B3 (e), or C1A-B12 (f). Interactions of CDR L3 residue 92 with the RBD are shown for C1A-B3 (g) or C1A-B12 (h). Interactions occurring at the base of CDR H1 near the framework regions are shown for C1A-B3 (I), C1A-C2 (j), or BD-236 (PDB: 7CHB)^8^ (k). The color scheme for the RBD and antibody Fab is the same as the one showed in Figure 2b. “Germline” indicates baseline interactions occurring when a given residue is not somatically mutated; mutations are otherwise listed on top of the panel.

As has been well described in antibody responses to influenza virus and HIV^20,21^, somatic hypermutation that are not at antibody/antigen interfaces can nevertheless substantially contribute to affinity gains by influencing CDR loop configuration and flexibility. The V_H_ A24V mutation is a pocket-filling mutation that, through hydrophobic interactions with the side chain of V_H_ F27, would rigidify CDR H1 may “pre-configure” it in a conformation that is compatible with RBD binding (Fig. 4i-j). V_H_ F27 is frequently mutated to a smaller hydrophobic residues during somatic hypermutation; it is replaced by an isoleucine in C1A-H5, BD-604, and BD-236^8^, by a leucine in CC12.1^12^, and by a valine in CV30^19^ (Fig. 4k; Supplementary Fig. 5a and 5c). In contrast to the V_H_ A24V mutation, replacing V_H_ F27 with smaller hydrophobic residue would likely make CDR H1 more flexible as opposed to rigidifying it, and this added flexibility could allow optimization of local polar contacts, particularly as additional mutations are introduced during affinity maturation (the T28I change in addition to the F27V mutation in CV30)^19^ (Supplementary Fig. 5c).

Affinity is not the only property that may be beneficial to an effective antibody response^22^, and antibody combining site diversity may provide broader protection against pathogens that are antigenically variable and evolve over time^23^. As examples of BCR diversification that could result in a loss of RBD affinity, the of V_H_ S56A mutation in C1A-B12 removes a polar contact with RBD D420, and the Y58F mutation in CC12.1 removes a polar contact with the backbone carbonyl of RBD T415 (Fig. 4f; Supplementary Fig. 5f-g).

IGHV3-53/3-66-derived SARS-CoV-2 neutralizing antibodies usually have short CDR H3 loops to avoid clashes with the RBD surface^12^ (Supplementary Fig. 4). Six of the seven clonally related IGH/V3-53 antibodies we isolated contain the S100aR mutation in CDR H3 with independent substitutions at the nucleotide level (Fig. 5a), suggesting that this adaptation was recurrently selected for during the affinity maturation process. We observed two alternate conformations for the R100a side chain in the C1A-B12 Fab/RBD structure; it can either contact the side chain of RBD Q493 or the backbone carbonyl of RBD S494 (Fig. 5b). The R100a side chain also helps position neighboring antibody residues to make additional contacts with the RBD as part of a larger network of polar interactions involving water molecules.

**Figure 5.**
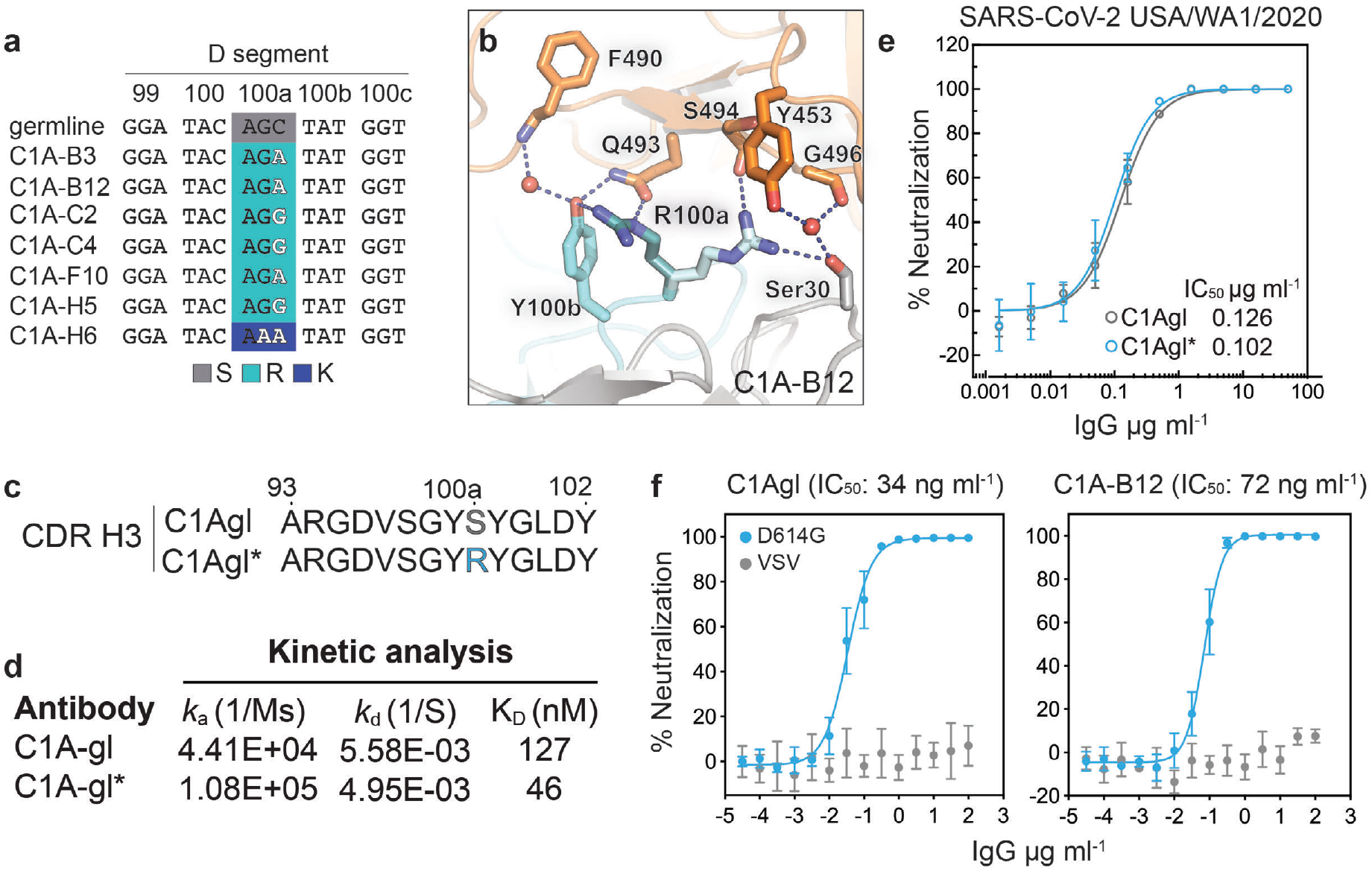
A germline revertant antibody neutralizes SARS-CoV-2. (a) Nucleotide sequences of D segment of C1A-IGHV3 antibodies. Substitutions acquired during somatic mutation causing the S100aR or S100aK substitutions are highlighted. (b) Ribbon diagram of C1A-B12/RBD complex showing interactions occurring with alternate side chain conformers of CDR H3 residue R100a. (c) Amino acid sequences for CDR H3 loops of germline revertant antibodies C1Agl and C1Agl*. (d) Results of kinetic analysis of binding for Fabs on immobilized SARS-CoV-2 RBD as measured by BLI. (e) Results of PRNT assay with infectious SARS-CoV-2 (strain USA/WA1/2020) and the indicated monoclonal antibodies. Data are normalized to a no antibody control. Means ± standard deviation from three experiments performed in triplicate (n=6). Error bars indicate standard deviation. For some data points, error bars are smaller than symbols. (f) Dose response neutralization assay results with SARS-CoV-2 lentivirus pseudotype with the D614G mutation. Data are normalized to a no antibody control. Means ± standard deviation from two experiments performed in triplicate (n=6). For some data points, error bars are smaller than symbols.

### Germline revertant antibodies neutralize SARS-CoV-2

To better understand the selective pressure driving the expansion of the IGVH3-53 class of SARS-CoV-2 neutralizing antibodies and the role of the S100aR somatic change in affinity maturation, we generated an antibody revertant in which all positions are reverted to their germline counterparts (C1A-gl), and another that only retains the S100aR substitution (C1A-gl*) (Fig. 3 and Fig. 5c). Monomeric C1A-gl and C1A-gl* Fabs bound the SARS-CoV-2 S RBD with affinities of 127 nM and C1A-gl* and 46 nM, respectively (Fig. 5d and Supplementary Fig. 3). The effect of the mutation was most pronounced on antibody on-rate, suggesting that the CDR H3 S100aR substitution allows the antibody to more effectively dock onto the RBD. C1A-gl and C1A-gl* IgG neutralized infectious SARS-CoV-2 with IC_50_ values of 0.126 and 0.102 μg ml^-1^, respectively (Fig. 4e).

We next sought to address whether C1A-IGHV3 antibodies could neutralize SARS-CoV-2 containing the D614G mutation in S, which is found in a circulating SARS-CoV-2 strain with increased infectivity ^24–26^. C1A-gl, which binds the RBD with 126 nM affinity, and C1A-B12, which binds the RBD with thirty-fold higher affinity (KD of 4 nM), neutralized SARS-CoV-2 S D614G pseudotypes with comparable IC_50_ values (74 and 34 ng ml^-1^, respectively) (Fig. 5f). Taken together, our results suggest that once a threshold RBD affinity is reached for IGHV3-53 antibodies, gains in affinity are not necessarily associated with more potent virus neutralization.

## Discussion

In the absence of effective vaccines or therapeutics, the SARS-CoV-2 pandemic has continued unabated with most of the world now facing new waves of infection despite aggressive infection control measures. A detailed understanding of the human antibody response to this novel virus will be required to design safe and effective agents to be used as prophylaxis or early treatment of SARS-CoV-2 infection, including countermeasures that anticipate changes in the virus that could lead to antibody neutralization escape. Multiple groups have now reported on IGHV3-53/3-66-derived antibodies isolated from COVID-19 convalescent donors that carry low rates of somatic mutation and potently neutralize SARS-CoV-2 – these recognize the RBD with essentially the same binding mode (Supplementary Fig. 4)^3,4,8,10-13^. In the donor we studied, the neutralizing antibody response was dominated by this class of antibodies (Supplementary Fig. 1 and Supplementary Table 1), suggesting that in certain scenarios, the preponderance of such antibodies could drive selective evolution of antibody neutralization escape.

The high “baseline affinity” IGHV3-53/3-66-derived germline antibodies have for the SARS-CoV-2 RBD is likely driven by the extensive network of reciprocal polar contacts these make with both main and side chain atoms of the RBD (Fig. 2, Fig. 3, and Supplementary Fig. 5). Our ability to detect robust RBD binding for monomeric Fabs representing germline revertant antibody sequences (Fig. 5) in an assay that does not take into account the avidity that would be observed with B cell receptors engaging trimeric S suggests that strong selective pressure drives the evolution of antibody responses against this epitope. Our results stand somewhat in contrast to those observed with antibody CV30, an IGHV3-53/IGVK3-20 antibody for which reversion of its only two substitutions (V_H_ F27V and T28I) with respect to the germline antibody, results in a change in affinity from 3.6 nM to 407 nM and in an almost sixty fold change in neutralization IC_50_ value (from 0.030 μg ml^-1^ to 16.5 μg ml^-1^)^19^. As described in our analysis, the V_H_ F27V and T28I mutations may respectively affect loop dynamicity and help optimize the geometry of CDR H1 contacts with the RBD^19^. The lack of a drastic change in affinity with reversion of germline antibody sequences with C1A-IGHV3-53 antibodies suggest that these take better advantage of antigen complementarity afforded by their CDR H3 loop and light chain gene (IGVK1-9 for C1A-IGHV3-53 antibodies and IGVK3-20 for CV30) (Supplementary Fig. 4).

While we did not observe a correlation between RBD affinity and virus neutralization, we did not measure affinities with full length SARS-CoV-2 S. IGHV3-53/3-66 neutralizing antibodies can only bind the RBD when it is “up”^8,16^, and efficient virus neutralization by engagement of an epitope that is transiently exposed likely requires optimization of binding on- and off-rates in addition to gains in overall affinity.

Mutations that allow SARS-CoV-2 S to escape neutralization by antibodies that compete with ACE2 binding have been observed *in vitro*^27^ and recently in circulating strains^28^. Although SARS-CoV-2 encodes an exonuclease that increases the fidelity of replication of its large RNA genome, recurrence of an identical antibody response in multiple COVID-19 convalescent individuals suggests that selective pressure against the RBD-binding site for ACE2 is substantial. To broadly protect against emerging S variants and to counter evolution of neutralization escape over time as the virus circulates in humans, vaccine design efforts may need to focus on potent neutralizing antibodies binding additional sites on SARS-CoV-2 S, rather than on clonal expansion of one or a limited set of IGVH3-53/3-66-derived antibodies, as occurred during natural infection of the convalescent donor we studied.

## Methods

### Donors

This study was approved by the Harvard Medical School Office of Human Research Administration Institutional Review Board (IRB20-0365) as was the use of healthy donor control blood (IRB19-0786). We received informed, written consent from a healthy adult male participant (C1) who recovered from confirmed SARS2-CoV-2 infection, with mild illness not requiring hospitalization, five weeks before blood donation. We isolated C1 and control donor PBMCs by Ficoll-Plaque (GE Healthcare) density centrifugation.

### Single B cell sorting and antibody cloning

We stained and sorted single memory B cells as previously described^29^ using a MoFlo Astrios EQ Cell Sorter (Beckman Coulter). Briefly, we enriched B cells by incubating PBMCs with anti-CD20 MicroBeads (Miltenyi Biotec) followed by magnetic separation on a MACS LS column (Miltenyi Biotec) according to the manufacturer’s instructions. We washed, counted, and resuspended the B cells in phosphate buffered saline (PBS) containing 2% (v/v) FBS. We adjusted the B cells to a density of 1×10^7^ cells and incubated cells with biotinylated SARS CoV-2 spike (S2P) at a concentration of 5 μg ml^-1^ on ice for 30 min. After washing three times and resuspending the cells, we added anti-IgG-APC antibody (BD Biosciences), anti-CD19-FITC antibody (BD Biosciences), and streptavidin-PE (Invitrogen). After incubating the cells on ice for 30 min, we washed cells three times in PBS containing 2% (v/v) FBS and passed the suspension through a cell strainer before sorting.

We performed single cell cDNA synthesis using SuperScript™ III reverse transcriptase (Invitrogen) followed by nested PCR amplification to obtain the IgH, Igλ, and IgK variable segments from memory B cells as previously described^30^. We used IMGT/V-QUEST^31^ (http://www.imgt.org) to analyze IgG gene usage and the extent of variable segment somatic hypermutation. The variable segments were cloned into the pVRC8400 vector for expression of the IgG and Fab constructs as previously described^32^.

### Cells and viruses

We maintained HEK293T cells (ATCC CRL-11268) in Dulbecco’s Modified Eagle’s Medium (DMEM) supplemented with 10% (v/v) fetal bovine serum (FBS) and 1% (v/v) penicillinstreptomycin and Expi293F™ cells (Thermo Fisher Scientific) in Expi293™ expression medium (Gibco) supplemented with 1% (v/v) penicillin-streptomycin. A HEK293T-hACE2 stable cell line was a gift from Huihui Mou and Michael Farzan and an Expi293F-His6-tagged SARS-CoV-2 S2P stable cell line that expresses a gift from Bing Chen. We maintained these in the same media with the addition of 1 μg ml^-1^ puromycin. We maintained HEK293T cells grown in suspension in FreeStyle 293 Expression Medium (Gibco) and HEK293S GnTI^-/-^ cells (ATCC CRL-3022) in Freestyle 293 Expression Medium supplemented with 2% ultra-low IgG FBS (Gibco). We maintained, was maintained in adherent culture with DMEM supplemented with 1% (v/v) GlutaMax (Gibco), 1% (v/v) penicillin-streptomycin, 10% (v/v) FBS and 1μg ml^-1^ puromycin. The cell line was then adapted to suspension culture and maintained in Expi293™ expression medium supplemented with 1% (v/v) penicillin-streptomycin and 1 μg ml^-1^ puromycin (Gibco).

Passage 4 SARS-CoV-2 USA/WA1/2020^33^ was received from the University of Texas Medical Branch. A T225 flask of VeroE6 cells was inoculated with 90 μl starting material in 15 ml DMEM containing 2% (v/v) of heat inactivated FBS (HI-FBS) and incubated in a humidified incubator at 37 °C with periodic rocking for 1 h. After 1 h, 60 ml of DMEM / 2% (v/v) HI-FBS was added without removing the inoculum and incubated again at 37 °C. The flask was observed daily for progression of cytopathic effect and stock was harvested at 66 h post-inoculation. Stock supernatant was harvested and clarified by centrifugation at 5,250 relative centrifugal field (RCF) at 4°C for 10 min and the HI-FBS concentration was increased to 10% (v/v) final concentration.

### Protein production

For single B cell sorting we cloned a construct for SARS-CoV-2 S (GenBank ID: QHD43416.1 residues 16-1208) with a “GSAS” substitution at the furin cleavage site (residues 682-685), stabilized in the prefusion conformation through proline substitutions at residues 986 and 987^14^, and a C-terminal foldon trimerization motif followed by a BirA ligase site, a Tobacco Etch Virus (TEV) protease site, a FLAG tag, and a His6-tag into a pHLsec vector^34^, which contains its own secretion signal sequence. We note that two N-terminal S residues (residues 14 and 15) downstream of the native S signal peptide were inadvertently omitted from the S2P construct during subcloning. We transfected Expi293F™ cells using an ExpiFectamine™ transfection kit (Thermo Fisher Scientific) according to the manufacturer’s protocol. We purified the protein using anti-FLAG M2 Affinity Gel (Sigma) according to manufacturer’s protocol and removed the FLAG tag and His6-tag with TEV digestion followed by reverse nickel affinity purification and size-exclusion chromatography on a Superose 6 Increase column (GE Healthcare Life Sciences). We biotinylated the protein with BirA ligase as previously described^32^.

To obtain recombinant S2P for ELISAs, we used Ni Sepharose^®^ Excel (GE Healthcare Life Sciences) to purify His6-tagged SARS-CoV-2 S2P from the supernatant of Expi293F cells stably expressing this protein. We further purified the protein using size exclusion chromatography on a Superpose 6 Increase column.

We expressed and purified recombinant monoclonal antibodies and Fab fragments using the pVRC8400 vector as previously described^32^. To generate the CR3022 control Fab, we amplified its variable heavy chain and light chain gene regions (GenBank IDs: DQ168569.1 and DQ168570.1) from cDNA (a gift from Galit Alter) and subcloned these into the pVRC8400 vector. We used size exclusion chromatography on a Superdex 200 Increase column (S200, GE Healthcare Life Sciences) for all Fabs, which eluted as single peaks at the expected retention volume.

We subcloned constructs for the SARS-CoV-2 S RBD (GenBank ID: QHD43416.1 residues 319-541) into the pHLsec^34^ vector for use in ELISAs, BLI binding studies, and X-ray crystallography. For ELISAs and crystallography the construct includes an N-terminal His6-tag, a TEV protease site and a short linker (amino acids SGSG). For BLI-binding assays, the construct includes an N-terminal His6-tag, followed by a TEV protease site, a BirA ligase site, and a 7-residue linker. We produced protein for ELISA and BLI-binding assays by using linear polyethylenimine (PEI) to transfect HEK293T cells grown in suspension and purified by nickel affinity purification. For BLI-binding assays the protein was digested with TEV protease to remove the His6-tag followed by reverse nickel affinity purification. We biotinylated protein with BirA ligase as previously described^35^, followed by a reverse nickel affinity purification step to remove BirA ligase, which contains a His6-tag and cannot be separated by size exclusion chromatography from the SARS-CoV-2 RBD due to its similar size. For crystallography, we produced the protein by PEI transfection of GnTI^-/-^ HEK293S cells grown in suspension or HEK293T cells grown in suspension and also in presence of kifunensine (5 μM), purified by nickel affinity purification, and removed the His6-tag by TEV digestion followed by reverse nickel affinity purification. As a final step, we used size exclusion on a S200 Increase column, in which each recombinant RBD protein ran as a single peak at the expected retention volume.

We subcloned the ectodomain of human ACE2 (GenBank ID: BAB40370.1 residues 18-740, with cDNA obtained as a gift cDNA from Michael Farzan, with a C-terminal Fc tag into a pVRC8400 vector containing human IgG1 Fc (a gift from Aaron Schmidt). We expressed the protein in Expi293F™ cells using an ExpiFectamine™ transfection kit according to the manufacturer’s protocol, and purified the protein using MabSelect SuRE Resin using the manufacturer’s protocol, followed by size exclusion chromatography on a Superose 6 Increase column, with the protein eluting at the expected retention volume.

### Crystallization

We prepared each Fab:SARS-CoV-2 RBD complex by mixing RBD with 1.5 molar excess of Fab. The mixtures were incubated at 4°C for 1 h prior to purification on a Superdex 200 Increase SEC column (GE Healthcare Life Sciences) in buffer containing 150mM NaCl, 25 mM Tris-HCl, pH 7.5. Each complex co-eluted as a single peak at expected retention volume. We adjusted the concentration of each complex to 13 mg ml^-1^ and screened for crystallization conditions in hanging drops containing 0.1 μl of protein and 0.1μl of mother liquor using a Mosquito protein crystallization robot (SPT Labtech) with commercially available screens (Hampton Research) (see Key Resources Table). Crystals grew within 24 h for the C1A-B12 Fab:RBD complex in 0.1 M BICINE pH 8.5, 20% (w/v) polyethylene glycol 10,000, for the C1A-B3 Fab:RBD complex in 0.2 M Ammonium phosphate dibasic, 20% (w/v) polyethylene glycol 3,350; for the C1A-C2 Fab:RBD complex in 0.03 M citric acid, 0.07M BIS-TRIS propane pH 7.6, 20% (w/v) polyethylene glycol 3,350, and for C1A-F10 Fab:RBD complex in 0.10% (w/v) n-Octyl-B-glucoside, 0.1 M Sodium citrate tribasic dihydrate pH 4.5, and 22% (w/v) polyethylene glycol 3,350.

### Structure determination

All crystals were flash frozen in mother liquor supplemented with 15% (v/v) glycerol as cryoprotectant. We collected single crystal X-ray diffraction data on Eiger X 16M pixel detectors (Dectris) at a wavelength of 0.979180 Å at the Advanced Photon Source (APS, Argonne, IL) NE-CAT beamline 24-ID-E for the C1A-B12 Fab:RBD and C1A-B3 Fab:RBD complexes and NE-CAT beamline 24-ID-C for the C1A-C2 Fab:RBD and C1A-F10 Fab:RBD complexes. Diffraction data were indexed and integrated using XDS (build 202 00131)^36^ and merged using AIMLESS (v0.5.32)^37^. The structure of C1A-B12 Fab:RBD (space group *P*2_1_2_1_2_1_) was determined by molecular replacement using Phaser (v2.8.3)^38^, with coordinates for the B38 Fab variable domain, constant domain and RBD (PDB ID: 7BZ5) (Wu et al., 2020) used as search models. Three copies were found in the asymmetric unit (ASU). We performed iterative model using O^39^ and refinement in Phenix (v1.18.2-3874)^40^ and Buster (v2.10.3)^41^, during which we also built alternative conformations where density was apparent. During refinements, we updated TLS groups calculated using Phenix^40^ and a python script, as well as occupancy restraints calculated in Buster. During model building, we also customized geometry restraints to prevent large displacement of unambiguous contacts in poor regions; the restraints were released once refinements became stable. Water molecules were automatically picked and updated in Buster, followed by manual examination and adjustment till late stage refinement. The structures of C1A-B3:RBD (space group *P*2_1_2_1_2_1_, 3 copies per ASU), C1A-C2:RBD (space group *C*222_1_, 1 copy per ASU) and C1A-F10:RBD (space group *C*222_1_, 1 copy per ASU) were determined using RBD and the C1A-B12 Fab variable and constant domains as search ensembles with CDR and flexible loops truncated, with iterative model building and refinement as described above. Data collection, processing and refinement statistics are summarized in Figure S2.

### Structural Analysis

We analyzed the structures and generated figures using PyMOL (Schrödinger).

### Lentivirus pseudotype production

The SARS-CoV-2 S protein (Genbank ID: QJR84873.1 residues 1-1246) with a modified cytoplasmic sequence that includes HIV gp41 residues (NRVRQGYS) replacing C-terminal residues 1247-1273 of the S protein (gift from Nir Hacohen) was subcloned into the pCAGGS expression vector. We used Gibson assembly to generate the S D614G mutant. A pCAGGS expressor plasmid for VSV G was previously described^42^. To package lentivirus, we cotransfected HEK293T cells using lipofectamine™ 3000 (Thermo Fisher Scientific) with an envelope gene-encoding pCAGGS vector, a packaging vector containing HIV Gag, Pol, Rev, and Tat (psPAX2, provided by Didier Trono, Addgene #12260), and a transfer vector containing GFP (lentiCas9-EGFP, a gift from Phillip Sharp and Feng Zhang, Addgene #63592)^43^ in which we deleted Cas9. After 18 h, we changed the supernatant to DMEM containing 2 % FBS (v/v). We harvested supernatants after 48 and 72 h, centrifuged at 3000 x g for 5 min, and filtered the supernatants through a 0.45 μm filter. To concentrate lentivirus pseudotypes, we layered the supernatant on top of a 10% (v/v) sucrose cushion in 50 mM Tris-HCl pH 7.5, 100 mM NaCl, 0.5 mM EDTA and spun samples at 10,000 x g for 4 h at 4 °C. We removed supernatants and resuspended virus pellets in Opti-MEM containing 5% (v/v) FBS and stored these at −80 °C.

### Pseudotype neutralization experiments

We purified polyclonal IgG from human plasma samples using Pierce™ Protein G Ultra Link™ Resin (Thermo Fisher Scientific) following the manufacturer’s protocol. We pre-incubated polyclonal serum IgG or monoclonal antibodies at a concentration with SARS-CoV-2 or VSV G lentivirus pseudotypes in the presence of 0.5 μg ml^-1^ of polybrene for 1 h minutes at 37 °C. Virus antibody mixtures were added to HEK293T-hACE2 with incubation on cells at 37 °C for 24 h, and the media replaced with DMEM containing 10% (v/v) FBS, 1% (v/v) penicillin-streptomycin (v/v), and 1 μg ml^-1^ puromycin. We determined the percent of GFP positive cells by FACS with an iQue Screener PLUS (Intellicyt) 48 h after initial infection. We calculated percent relative entry by using the following equation: Relative Entry (%) = (% GFP positive cells in antibody well)/(%GFP positive cells in no antibody control). We calculated percent neutralization using the following equation: Neutralization (%) = 1 – (% GFP positive cells in nanobody well)/(% GFP positive cells in PBS alone well).

### Live virus PRNT experiments

Monoclonal antibody samples were serially diluted in Dulbecco’s Phosphate-Buffered Saline (DPBS, Gibco) using half-log dilutions starting at a concentration of 50 μg ml^-1^. Dilutions were prepared in triplicate for each sample and plated in triplicate. Each dilution was incubated at 37 °C for 1 h with 1,000 plaque-forming units ml^-1^ (PFU ml^-1^) of SARS-CoV-2 (isolate USA-WA1/2020). 200 μl of each dilution was added to the confluent monolayers of NR-596 Vero E6 cells (ATCC) in triplicate and incubated in a 5% CO_2_ incubator at 37 °C for 1 hour. The cells were rocked gently every 15 min to prevent monolayer drying. Cells were then overlaid with a 1:1 solution of 2.5% (v/v) Avicel^®^ RC-591 microcrystalline cellulose and carboxymethylcellulose sodium (DuPont Nutrition & Biosciences) and 2x Modified Eagle Medium (MEM – Temin’s modification, Gibco) supplemented with 100 X antibiotic-antimycotic (Gibco) and 100X GlutaMAX (Gibco) both to a final concentration of 2X, and 10% (v/v) FBS (Gibco). The plates were then incubated at 37 °C for 2 d. After two days, the monolayers were fixed with 10% (v/v) neutral buffered formalin for at least 6 h (NBF, Sigma-Aldrich) and stained with 0.2% (v/v) aqueous Gentian Violet (RICCA Chemicals) in 10% (v/v) neutral buffered formalin for 30 min, followed by rinsing and plaque counting.

### ELISA experiments

We coated NUNC Maxisorp plates (Thermo Fisher Scientific) with His6-tagged SARS-CoV-2 S2P, SARS-CoV-2 RBD, or LUJV GP1 in PBS overnight at 4 °C, followed by a blocking step with PBS containing 3% (v/v) BSA 0.02% (v/v) Tween. We incubated monoclonal antibodies at a concentration of 100 μg ml^-1^ for one hour. We detected bound antibody with horseradish peroxidase (HRP)-coupled anti-human (Fc) antibody (Sigma Aldrich catalog number A0170).

### Biolayer interferometry assays

We performed BLI experiments with an Octet RED96e (Sartorius). For affinity measurements, biotinylated SARS-CoV-2 RBD was loaded onto a streptavidin (SA) sensor (ForteBio) at 1.5 μg ml^-1^ in kinetic buffer (PBS containing 0.02% Tween and 0.1% BSA) for 100 s. After a baseline measurement for 60 s in kinetic buffer, antibody Fabs were associated for 300 s followed by a 300 s dissociation step.

For ACE2-Fc competition experiments, we loaded biotinylated SARS-CoV-2 RBD onto SA sensors (ForteBio) at 1.5 μg ml^-1^ for 80 s. We associated C1A-B12 Fab or CR3022-Fab at 250 nM or buffer for 180 s followed by an association with ACE2-Fc or CR3022 Fab at a concentration of 250 nM for 180 s. We allowed complexes to dissociate for 180 s.

## Author contributions

Conceptualization, S.A.C, L.E.C, J.P., A.C.; Investigation, S.A.C., L.E.C., J.P., A.C., L.G.A.M., S.S., R.I.J., A.G., J.A.; Writing, original draft: J.A.; Writing – Review and editing, S.A.C., L.E.C., J.P., A.C., L.G.A.M., S.S., R.I.J., A.G., J.A.; Funding acquisition, JA.

## Acknowledgements

This work is based upon research conducted at the Northeastern Collaborative Access Team (NE-CAT) beamlines, which are funded by the National Institute of General Medical Sciences from the National Institutes of Health (P30 GM124165). The Pilatus 6M detector on 24-ID-C beam line is funded by a NIH-ORIP HEI grant (S10 RR029205). This research used resources of the Advanced Photon Source, a U.S. Department of Energy (DOE) Office of Science User Facility operated for the DOE Office of Science by Argonne National Laboratory under Contract No. DE-AC02-06CH11357. We thank the staff at NE-CAT for assistance with X-ray data collection. Funding was provided by the Massachusetts Consortium on Pathogen Readiness (MassCPR) and the China Evergrande Group (J.A.). We thank Bing Chen for providing a SARS-CoV-2 S2P expressing stable cell line and Michael Farzan and Huihui Mou for providing HEK293T-hACE2 cells. We also thank Lindsey Baden, Michael Mina, and Brendan Blumenstiel for help with sample acquisition and SARS-CoV-2 testing.

## Data availability

All relevant data are available from the authors upon request. Protein Data Bank (PBD) identification numbers for the C1A-B3/RBD, C1A-F10/RBD, C1A-C2/RBD, and C1A-B12 RBD complexes are 7KFW, 7KFY, 7KFX, and 7KFV, respectively.

---

**Supplementary Figure 1.**
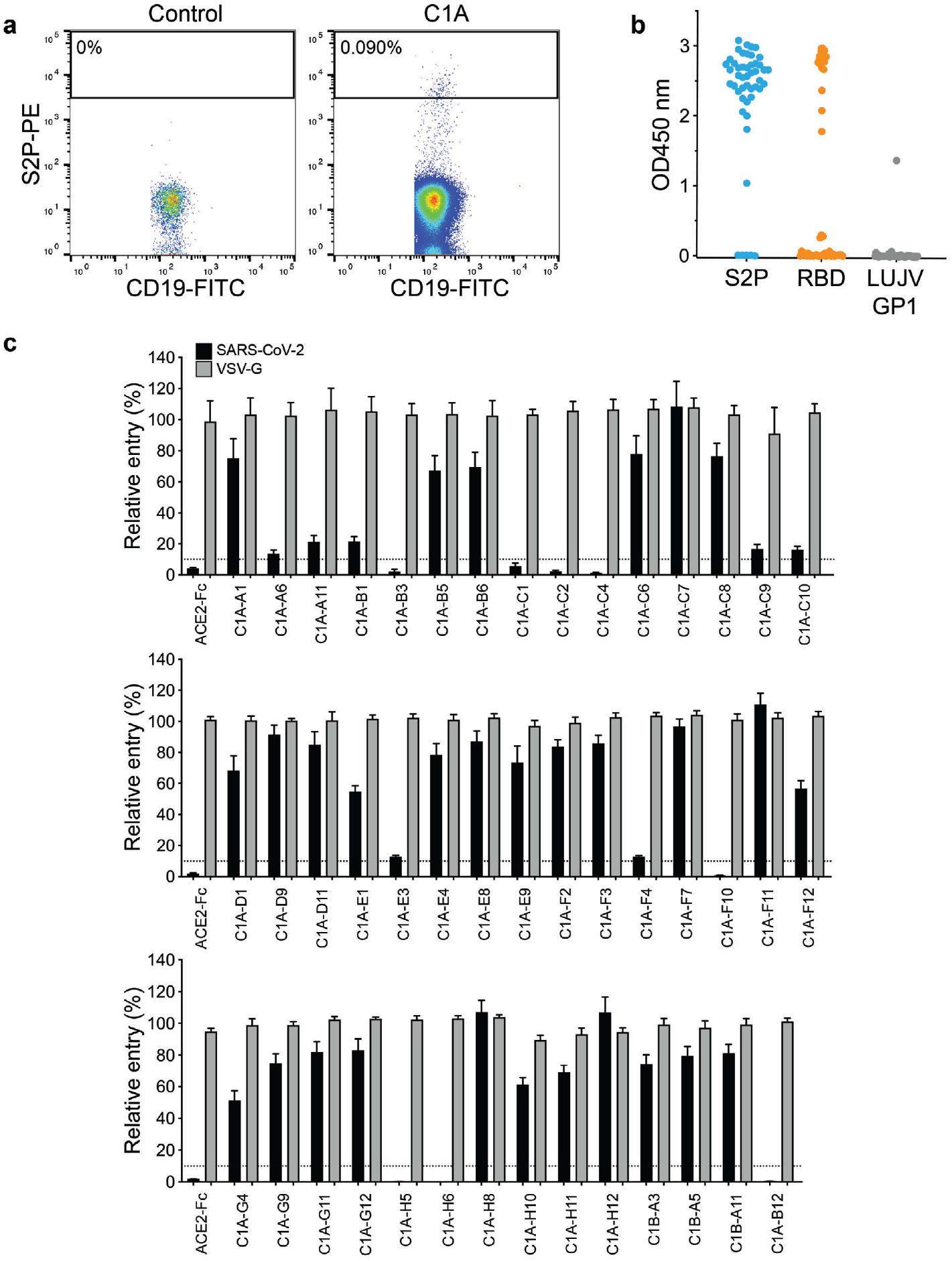
Monoclonal antibody isolation from a COVID-19 convalescent individual. (a) Density plot from a FACS experiment to isolate memory B cells that bind phycoerythrin (PE)-labelled streptavidin tetramers coupled to a prefusion-stabilized SARS-CoV-2 S construct (S2P-PE). The approximate location of the sorting gate is shown as a box, and the percentage of cells that fall within the gate is indicated. The left panel is for a control donor and the right panel is for a COVID-19 convalescent donor. CD19 is a B-cell marker. (b) Whisker plot showing ELISA values for IgG binding to S2P, the SARS-CoV-2 RBD, or the control protein Lujo virus (LUJV) GP1. Antibodies were added at a single concentration of 100 μg ml^-1^. (c) SARS-CoV-2 lentivirus pseudotypes were preincubated with 100 μg ml^-1^ of the indicated IgG or ACE2-Fc fusion protein (ACE2-Fc) and the mixture was used to infect HEK293T-hACE2 cells. Entry levels were quantified 48 h later using FACS. VSV lentivirus pseudotype is included as a control. Data are normalized to a no antibody control. Dashed line indicates 10% relative entry. Means ± standard deviation from two experiments performed in triplicate (n=6).

**Supplementary Figure 2.**
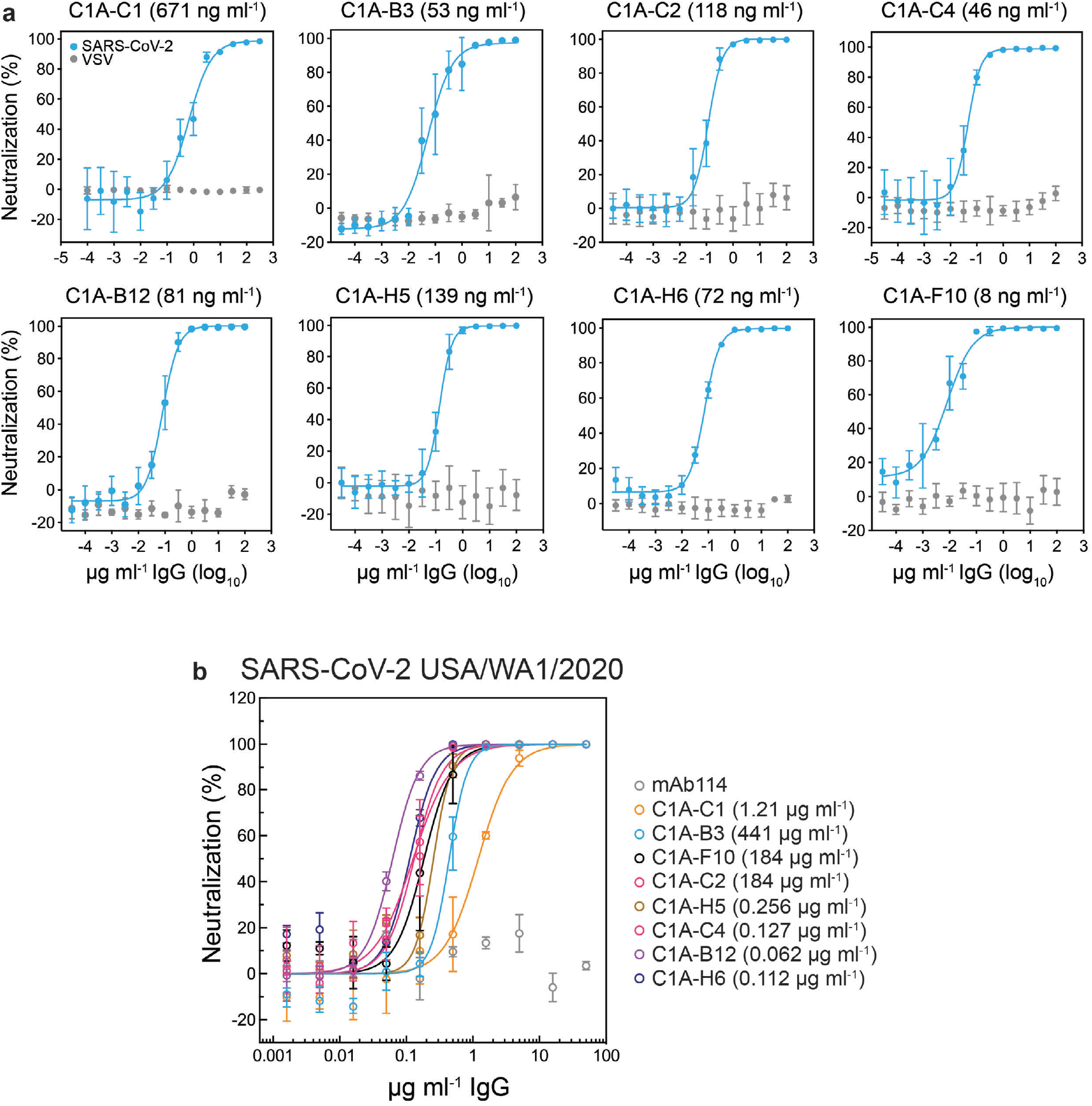
SARS-CoV-2 pseudotype and infectious virus neutralization assays. (a) SARS-CoV-2 lentivirus pseudotypes were pre-incubated with monoclonal antibodies at the indicated concentrations and the mixture was used to infect HEK293T-hACE2 cells. Entry levels were quantified 48 h later using FACS. VSV pseudotypes are included as a control. Data are normalized to a no antibody control. Means ± standard deviation from two experiments performed in triplicate (n=6). IC_50_ values are shown in parentheses. (b) Infectious SARS-CoV-2 (strain USA/WA1/2020) was incubated with monoclonal antibodies at the indicated concentration with infection of Vero E6 cells subsequently measured in PRNT assay^46^. Monoclonal antibody mAb114 is included as a control. Each monoclonal antibody was serially diluted in Dulbecco’s phosphate Buffered Saline (DPBS) using half-log dilutions starting at a concentration of 50 μg ml^-1^. Means ± standard deviation from three experiments performed in triplicate (n=9). Data are normalized to a no antibody control. For some data points, error bars are smaller than symbols.

**Supplementary Figure 3.**
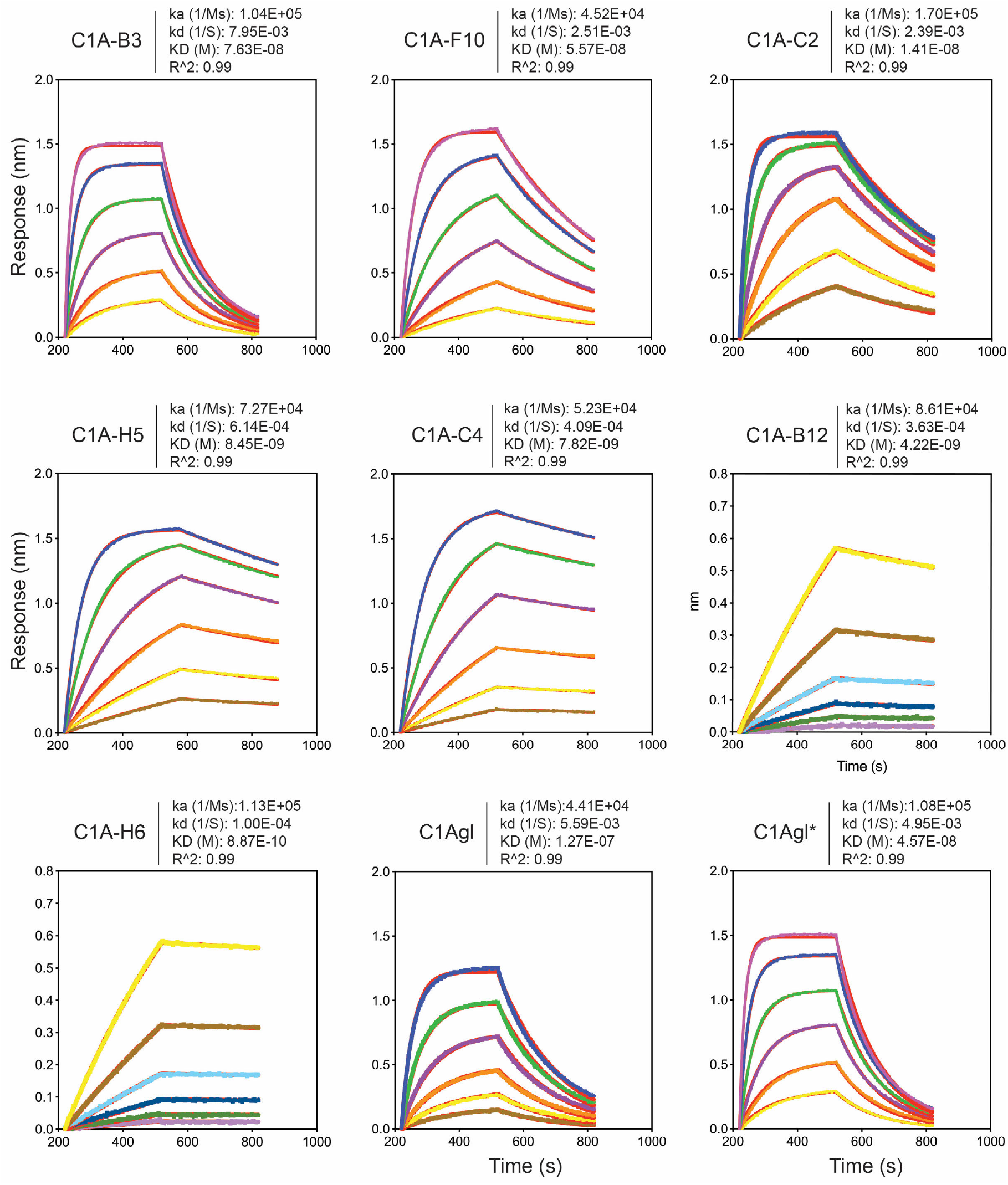
Fab binding kinetics to the SARS-CoV-2 receptor-binding domain. Fab affinities against the SARS-CoV-2 RBD were measured using biolayer interferometry (BLI). Red lines represent the fit for a 1:1 binding model, and alternate colors represent response curves measured at varying concentrations. Binding kinetics were measured for six concentrations of Fab at 2-fold dilution ranging from 500 to 15.6 nM (for Fab C1A-B3, C1A-F10, C1Agl, C1Agl*), 250 to 7.8 nM (C1A-C2, C1A-H5, C1A-C4), and from 15.6 to 0.49 nM (Fab C1A-B12 and C1A-H6), ensuring that each dilution series had concentrations both above and below the dissociation constant (KD). Representative results of two replicates for each experiment are shown.

**Supplementary Figure 4.**
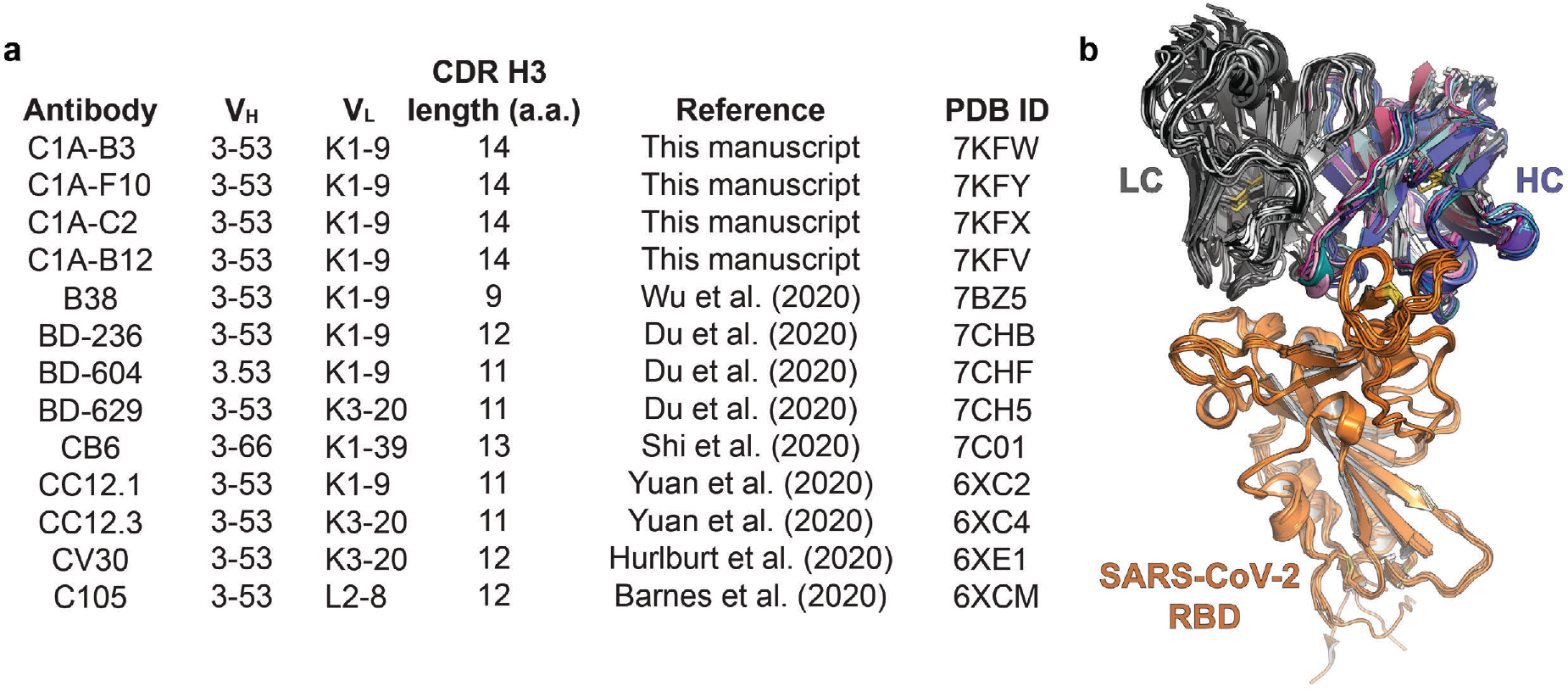
Structural comparison of IGVH3-53/3-66-derived antibodies. (a) Gene usage and CDR H3 lengths IGVH3-53/3-66 antibodies included in our analysis. CDR H3 length was determined using IMGT/V-QUEST definitions^31^. (b) Structural alignment of variable heavy (V_H_) and variable light (V_L_) portion of Fabs derived from IGHV3-53/3-66 bound to the SARS-CoV-2 RBD for all antibodies listed in (a). a.a.: amino acids. PDB ID: protein data bank identification code.

**Supplementary Figure 5.**
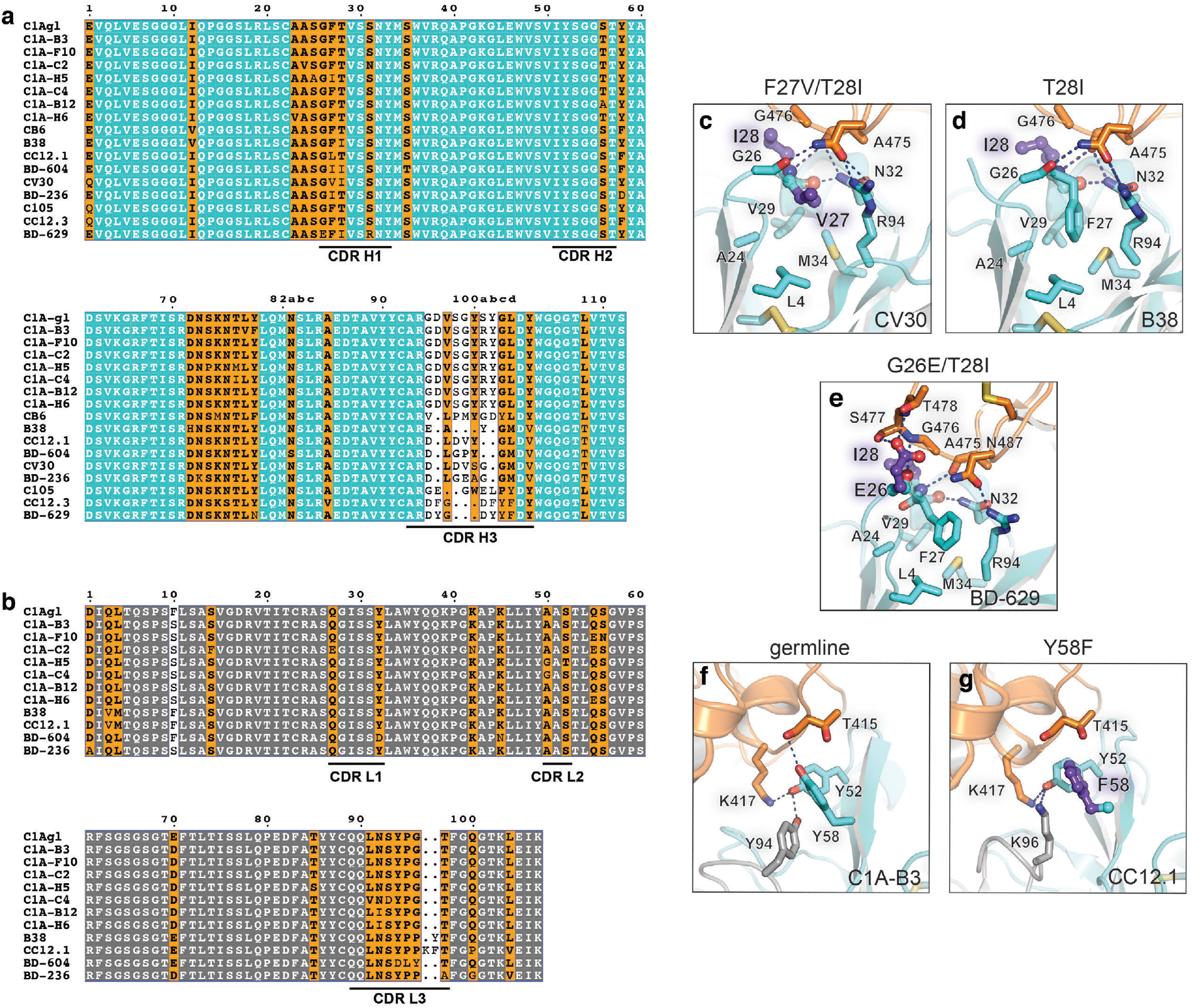
Sequence alignment with other reported IGHV3-53/3-66-derived antibodies. (a) Alignment of variable heavy chain and (b) IGLVK1-9-derived light chain sequences of IGHV3-53/3-66 antibody genes reported here and elsewhere. Antibody sequences were obtained from the RCSB record and protein data bank (PDB) IDs listed in Fig S4A. Panels were generated using ESPrit3^45^ and modified. The Kabat numbering scheme is used. (c) CV30-Fab/RBD complex (PDB: 6XE1^19^ showing interactions occurring with CDR H1 mutations F27V and T28I. (d) B38 Fab/RBD complex (PDB: 7BZ5)^4^ showing interactions occurring with the CDR H1 T28I mutation. (e) BD-629 Fab/RBD complex (PDB: 7CH5)^8^ showing interactions occurring with the CDR H1 G26E and T28I mutations. (f) C1A-B3 Fab/RBD complex showing interactions occurring with the germline CDR H2 residue Y’2. (g) CC12.1 Fab/RBD complex (PDB: 6XC2)^12^ showing interactions with the Y58F mutation.

**Supplementary Table 1.** Properties of monoclonal antibodies isolated from a COVID-19 convalescent individual.

| Andibody | V<sub>H</sub> gene | CDR H3 (a.a.) | Identity (%) | V<sub>L</sub> gene | CDR L3 (a.a.) | Identity (%) | Neut. | ELISA (OD 450 nm) |  |  |
| --- | --- | --- | --- | --- | --- | --- | --- | --- | --- | --- |
|  |  |  |  |  |  |  |  | S2P | RBD | Ctrl |
| C1A-A1 | IGHV1-69 | 15 | 99.31 | IGKV3-11 | 11 | 99.64 | - | 2.46 | 0.26 | 0.00 |
| C1A-A6 | IGHV3-11 | 15 | 98.96 | IGKV1-13 | 9 | 98.92 | + | 2.55 | 2.81 | 0.00 |
| C1A-A11 | IGHV3-30 | 20 | 99.31 | IGKV1-33 | 9 | 99.28 | + | 2.87 | 2.85 | -0.01 |
| C1A-B1 | IGHV1-2*02 | 18 | 97.57 | IGKV3-20 | 10 | 98.58 | + | 2.44 | 0.04 | 0.00 |
| C1A-B3 | IGHV3-53 | 14 | 97.89 | IGKV1-9 | 9 | 98.92 | ++ | 2.66 | 2.94 | 0.00 |
| C1A-B5 | IGHV4-59*11 | 11 | 94.74 | IGKV4-1*01 | 9 | 95.96 | - | 2.73 | 0.04 | 0.00 |
| C1A-B6 | IGHV1-24 | 14 | 98.61 | IGLV2-8 | 10 | 99.31 | - | 3.08 | 0.00 | 0.00 |
| C1A-B12 | IGHV3-53 | 14 | 98.6 | IGKV1-9 | 9 | 98.57 | ++ | 2.69 | 2.97 | -0.01 |
| C1A-C1 | IGHV3-23 | 14 | 99.31 | IGKV1-39 | 10 | 100 | ++ | 2.95 | 2.97 | -0.01 |
| C1A-C2 | IGHV3-53 | 14 | 98.6 | IGKV1-9 | 9 | 97.49 | ++ | 2.81 | 2.86 | -0.01 |
| C1A-C4 | IGHV3-53 | 14 | 98.25 | IGKV1-9 | 9 | 97.13 | ++ | 2.89 | 2.85 | 0.00 |
| C1A-C6 | IGHV1-69 | 15 | 98.96 | IGKV3-11 | 11 | 98.92 | - | 2.58 | 0.02 | 0.01 |
| C1A-C7 | IGHV4-39 | 12 | 99.31 | IGLV2-14 | 10 | 94.79 | - | 1.04 | 2.08 | 1.37 |
| C1A-C8 | IGHV1-24 | 16 | 99.65 | IGLV3-21 | 12 | 98.57 | - | 3.02 | 0.00 | -0.01 |
| C1A-C9 | IGHV1-69 | 14 | 97.92 | IGKV1-39 | 8 | 98.92 | + | 2.99 | 2.93 | 0.00 |
| C1A-C10 | IGHV3-23 | 17 | 98.61 | IGLV2-23 | 12 | 99.65 | + | 2.75 | 0.04 | 0.01 |
| C1A-D1 | IGHV4-39 | 18 | 91.75 | IGKV1-33 | 9 | 93.19 | - | 2.06 | 0.30 | 0.05 |
| C1A-D9 | IGHV3-30 | 20 | 98.61 | IGKV2-30 | 10 | 100 | - | 2.66 | 1.78 | 0.02 |
| C1A-D11 | IGHV1-69 | 14 | 98.26 | IGKV3-11 | 11 | 99.28 | - | 2.39 | 0.00 | -0.01 |
| C1A-E1 | IGHV3-9*01 | 13 | 89.58 | IGKV1-39 | 10 | 91.04 | - | 2.70 | 2.82 | -0.01 |
| C1A-E3 | IGHV1-18 | 16 | 98.61 | IGKV1-39 | 5 | 98.92 | + | 2.74 | 2.86 | 0.00 |
| C1A-E4 | IGHV3-30 | 14 | 97.92 | IGKV3-20 | 9 | 100 | - | 2.43 | 0.03 | 0.00 |
| C1A-E8 | IGHV3-48 | 21 | 97.22 | IGKV2-28 | 9 | 100 | - | 2.73 | 0.02 | 0.01 |
| C1A-E9 | IGHV3-30 | 14 | 100 | IGKV3-20 | 9 | 100 | - | 1.81 | 0.00 | 0.00 |
| C1A-F2 | IGHV3-30-3 | 15 | 99.65 | IGKV2-28 | 8 | 99.66 | - | 2.25 | 0.01 | 0.00 |
| C1A-F3 | IGHV4-61 | 13 | 98.63 | IGKV1-39 | 10 | 99.28 | - | 2.40 | 0.07 | 0.01 |
| C1A-F4 | IGHV3-30-3 | 14 | 99.65 | IGKV1-5 | 9 | 99.64 | + | 2.84 | 2.77 | 0.00 |
| C1A-F7 | IGHV3-15 | 11 | 91.84 | IGKV4-1 | 9 | 93.6 | - | 2.42 | 0.01 | 0.00 |
| C1A-F10 | IGHV3-53 | 14 | 97.89 | IGKV1-9 | 9 | 98.21 | ++ | 2.98 | 2.69 | 0.00 |
| C1A-F11 | IGHV1-69 | 21 | 89.24 | IGKV1-33 | 8 | 94.98 | - | 2.39 | 2.37 | 0.00 |
| C1A-F12 | IGHV4-39 | 15 | 90.03 | IGLV1-40 | 11 | 95.49 | - | 2.62 | 2.78 | 0.01 |
| C1A-G4 | IGHV3-23 | 17 | 99.65 | IGLV2-23 | 11 | 99.65 | - | 2.57 | 0.00 | 0.00 |
| C1A-G9 | IGHV3-30-3 | 14 | 98.61 | IGKV3-20 | 8 | 100 | - | 2.46 | 0.01 | 0.00 |
| C1A-G11 | IGHV1-69 | 12 | 97.92 | IGKV1-5 | 9 | 99.28 | - | 2.75 | 0.01 | 0.00 |
| C1A-G12 | IGHV3-30-3 | 14 | 98.26 | IGKV3-15 | 8 | 98.92 | - | 2.51 | 0.07 | 0.07 |
| C1A-H5 | IGHV3-53 | 14 | 97.54 | IGKV1-9 | 9 | 97.49 | ++ | 2.69 | 2.67 | 0.00 |
| C1A-H6 | IGHV3-53 | 14 | 98.95 | IGKV1-9 | 9 | 98.21 | ++ | 2.90 | 2.74 | 0.04 |
| C1A-H10 | IGHV3-21 | 18 | 99.31 | IGKV3-15 | 10 | 100 | - | 2.64 | 0.03 | 0.02 |
| C1A-H11 | IGHV3-30 | 16 | 98.61 | IGLV2-14 | 10 | 99.31 | - | 2.27 | 0.00 | 0.00 |
| C1A-H12 | IGHV4-39 | 15 | 98.28 | IGKV3-15 | 9 | 99.28 | - | 2.66 | 0.01 | 0.01 |
| C1B-A3 | IGHV3-30-3 | 14 | 99.65 | IGKV3-11 | 9 | 100 | - | 2.00 | 0.02 | 0.01 |
| C1B-A5 | IGHV3-30-3 | 14 | 99.65 | IGKV3-20 | 9 | 98.94 | - | 2.20 | 0.01 | 0.01 |
| C1B-A11 | IGHV3-30-3 | 14 | 98.26 | IGKV3-20 | 9 | 99.29 | - | 2.35 | 0.28 | 0.00 |

Antibodies highlighted in gray are somatic variants of the same antibody. CDR loop lengths are shown as numbers of amino acids (a.a.). ELISA values are colored in shades of blue according to their magnitude; darker shades are reflective of a stronger signal.

**Supplementary Table 2.** Data collection and refinement statistics

|  | C1A-B3/RBD<sup>a</sup><br>(PDB 7KFV) | C1A-B12/RBD<sup>a</sup><br>(PDB 7KFW) | C1A-C2/RBD<sup>a</sup><br>(PDB 7KFX) | C1A-F10/RBD<sup>a</sup><br>(PDB 7KFY) |
| --- | --- | --- | --- | --- |
| **Data collection** |  |  |  |  |
| Space group | *P*2<sub>1</sub>2<sub>1</sub>2<sub>1</sub> | *P*2<sub>1</sub>2<sub>1</sub>2<sub>1</sub> | *C*222<sub>1</sub> | *C*222<sub>1</sub> |
| Cell dimensions |  |  |  |  |
| *a*, *b*, *c* (Å) | 84.8, 113.3,<br>268.89 | 84.8, 113.3,<br>268.9 | 83.5, 149.2,<br>146.1 | 85.7, 146.8,<br>144.6 |
| $\alpha$, $\beta$, $\gamma$ (°) | 90.00, 90.00,<br>90.00 | 90.00, 90.00,<br>90.00 | 90.00, 90.00,<br>90.00 | 90.00, 90.00,<br>90.00 |
| Resolution (Å) | 200-2.77 (2.94-2.77)<sup>b</sup> | 200-2.10 (2.23-2.10)<sup>b</sup> | 200-2.22 (2.36-2.22)<sup>b</sup> | 200-2.16 (2.29-2.16)<sup>b</sup> |
| *R*<sub>sym</sub> | 0.38 (3.17)<sup>b</sup> | 0.238 (2.01)<sup>b</sup> | 0.136 (2.23)<sup>b</sup> | 0.118 (1.13)<sup>b</sup> |
| *R*<sub>pim</sub> | 0.11 (1.03)<sup>b,c</sup> | 0.095 (0.85)<sup>b,c</sup> | 0.062 (1.01)<sup>b,c</sup> | 0.076 (0.61)<sup>b,c</sup> |
| *I* / $\sigma$ | 5.4 (0.7)<sup>b</sup> | 9.0 (1.4)<sup>b</sup> | 6.9 (0.6)<sup>b</sup> | 6.2 (0.7)<sup>b</sup> |
| Completeness (%) | 98.1 (88.6)<sup>b</sup> | 98.9 (94.7)<sup>b</sup> | 99.1 (97.1)<sup>b</sup> | 94.0 (89.4)<sup>b</sup> |
| Redundancy | 6.9 (6.5)<sup>b</sup> | 7.0 (6.4)<sup>b</sup> | 3.6 (3.6)<sup>b</sup> | 2.1 (2.0)<sup>b</sup> |
| **Refinement** |  |  |  |  |
| Resolution (Å) | 133.92-2.79 | 134.44-2.10 | 74.60-2.23 | 74.04-2.16 |
| No. reflections | 63645 | 149225 | 44541 | 48373 |
| *R*<sub>work</sub> / *R*<sub>free</sub> | 0.19/0.23 | 0.18/0.22 | 0.19/0.23 | 0.19/0.22 |
| No. atoms |  |  |  |  |
| Protein | 14414 | 14511 | 4861 | 4855 |
| Ligand/ion | 42 | 42 | 14 | 14 |
| Water | 400 | 1642 | 325 | 532 |
| *B*-factors (Å<sup>2</sup>) |  |  |  |  |
| Protein | 88 | 47 | 63 | 52 |
| Ligand/ion | 96 | 99 | 103 | 101 |
| Water | 66 | 53 | 65 | 55 |
| R.m.s. deviations |  |  |  |  |
| Bond lengths<br>(Å) | 0.008 | 0.008 | 0.008 | 0.008 |
| Bond angles (°) | 0.980 | 0.970 | 1.000 | 1.000 |

<sup>a</sup> Numbers of crystals for C1A-B3, C1A-B12, C1A-C2 and C1A-F10 data were 1 each.
<sup>b</sup> Values in parentheses are for the highest-resolution shell.
<sup>c</sup> Values from program *aimless*.

## References

1 Zhou, P. et al. A pneumonia outbreak associated with a new coronavirus of probable bat origin. Nature, 270–273, doi:10.1038/s41586-020-2012-7 (2020).

2 Hoffmann, M. et al. SARS-CoV-2 Cell Entry Depends on ACE2 and TMPRSS2 and Is Blocked by a Clinically Proven Protease Inhibitor. Cell, 271–280.e278 doi:10.1016/j.cell.2020.02.052 (2020).

3 Robbiani, D. F. et al. Convergent antibody responses to SARS-CoV-2 in convalescent individuals. Nature, 437–442, doi:10.1038/s41586-020-2456-9 (2020).

4 Wu, Y. et al. A noncompeting pair of human neutralizing antibodies block COVID-19 virus binding to its receptor ACE2. Science, 1274–1278, doi:10.1126/science.abc2241 (2020).

5 Hansen, J. et al. Studies in humanized mice and convalescent humans yield a SARS-CoV-2 antibody cocktail. Science, 1010–1014, doi:10.1126/science.abd0827 (2020).

6 Chi, X. et al. A neutralizing human antibody binds to the N-terminal domain of the Spike protein of SARS-CoV-2. Science 369, 650–655, doi:10.1126/science.abc6952 (2020).

7 Liu, L. et al. Potent neutralizing antibodies against multiple epitopes on SARS-CoV-2 spike. Nature, 450–456, doi:10.1038/s41586-020-2571-7 (2020).

8 Du, S. et al. Structurally Resolved SARS-CoV-2 Antibody Shows High Efficacy in Severely Infected Hamsters and Provides a Potent Cocktail Pairing Strategy. Cell, doi:10.1016/j.cell.2020.09.035 (2020).

9 Lefranc, M.-P. & Lefranc, G. The Immunoglobulin FactsBook. (2014).

10 Shi, R. et al. A human neutralizing antibody targets the receptor-binding site of SARS-CoV-2. Nature 584, 120–124, doi:10.1038/s41586-020-2381-y (2020).

11 Rogers, T. F. et al. Isolation of potent SARS-CoV-2 neutralizing antibodies and protection from disease in a small animal model. Science, 956–963, doi:10.1126/science.abc7520 (2020).

12 Yuan, M. et al. Structural basis of a shared antibody response to SARS-CoV-2. Science, 1119–1123, doi:10.1126/science.abd2321 (2020).

13 Seydoux, E. et al. Analysis of a SARS-CoV-2-Infected Individual Reveals Development of Potent Neutralizing Antibodies with Limited Somatic Mutation. Immunity 53, 98–105 e105, doi:10.1016/j.immuni.2020.06.001 (2020).

14 Wrapp, D. et al. Cryo-EM structure of the 2019-nCoV spike in the prefusion conformation. Science 367, 1260–1263, doi:10.1126/science.abb2507 (2020).

15 Walls, A. C. et al. Structure, Function, and Antigenicity of the SARS-CoV-2 Spike Glycoprotein. Cell, 281–292.e286, doi:10.1016/j.cell.2020.02.058 (2020).

16 Barnes, C. O. et al. Structures of Human Antibodies Bound to SARS-CoV-2 Spike Reveal Common Epitopes and Recurrent Features of Antibodies. Cell 182, 828–842.e816, doi:10.1016/j.cell.2020.06.025 (2020).

17 Barnes, C. O. et al. SARS-CoV-2 neutralizing antibody structures inform therapeutic strategies. Nature, doi:10.1038/s41586-020-2852-1 (2020).

18 Yuan, M. et al. A highly conserved cryptic epitope in the receptor binding domains of SARS-CoV-2 and SARS-COV. Science 368, 630–633, doi:10.1126/science.abb7269 (2020).

19 Hurlburt, N. K. et al. Structural basis for potent neutralization of SARS-CoV-2 and role of antibody affinity maturation. bioRxiv, 2020.2006.2012.148692, doi:10.1101/2020.06.12.148692 (2020).

20 Schmidt, A. G. et al. Preconfiguration of the antigen-binding site during affinity maturation of a broadly neutralizing influenza virus antibody. Proc Natl Acad Sci U S A 110, 264–269, doi:10.1073/pnas.1218256109 (2013).

21 Klein, F. et al. Somatic mutations of the immunoglobulin framework are generally required for broad and potent HIV-1 neutralization. Cell 153, 126–138, doi:10.1016/j.cell.2013.03.018 (2013).

22 Eisen, H. N. Affinity enhancement of antibodies: how low-affinity antibodies produced early in immune responses are followed by high-affinity antibodies later and in memory B-cell responses. Cancer Immunol Res 2, 381–392, doi:10.1158/2326-6066.CIR-14-0029 (2014).

23 McCarthy, K. R., Raymond, D. D., Do, K. T., Schmidt, A. G. & Harrison, S. C. Affinity maturation in a human humoral response to influenza hemagglutinin. Proc Natl Acad Sci U S A, doi:10.1073/pnas.1915620116 (2019).

24 Korber, B. et al. Tracking Changes in SARS-CoV-2 Spike: Evidence that D614G Increases Infectivity of the COVID-19 Virus. Cell 182, 812–827.e819, doi:10.1016/j.cell.2020.06.043 (2020).

25 Zhang, L. et al. The D614G mutation in the SARS-CoV-2 spike protein reduces S1 shedding and increases infectivity. bioRxiv, doi:10.1101/2020.06.12.148726 (2020).

26 Yurkovetskiy, L. et al. Structural and Functional Analysis of the D614G SARS-CoV-2 Spike Protein Variant. Cell, 739–751.e738, doi:10.1016/j.cell.2020.09.032 (2020).

27 Baum, A. et al. Antibody cocktail to SARS-CoV-2 spike protein prevents rapid mutational escape seen with individual antibodies. Science, 1014–1018, doi:10.1126/science.abd0831 (2020).

28 Thomson, E. C. et al. The circulating SARS-CoV-2 spike variant N439K maintains fitness while evading antibody-mediated immunity. bioRxiv, 2020.2011.2004.355842, doi:10.1101/2020.11.04.355842 (2020).

29 Scheid, J. F. et al. A method for identification of HIV gp140 binding memory B cells in human blood. J Immunol Methods 343, 65–67, doi:10.1016/j.jim.2008.11.012 (2009).

30 Scheid, J. F. et al. Sequence and structural convergence of broad and potent HIV antibodies that mimic CD4 binding. Science 333, 1633–1637, doi:10.1126/science.1207227 (2011).

31 Brochet, X., Lefranc, M. P. & Giudicelli, V. IMGT/V-QUEST: the highly customized and integrated system for IG and TR standardized V-J and V-D-J sequence analysis. Nucleic Acids Res 36, W503–508, doi:10.1093/nar/gkn316 (2008).

32 Clark, L. E. et al. Vaccine-elicited receptor-binding site antibodies neutralize two New World hemorrhagic fever arenaviruses. Nat Commun 9, 1884, doi:10.1038/s41467-018-04271-z (2018).

33 Harcourt, J. et al. Severe Acute Respiratory Syndrome Coronavirus 2 from Patient with Coronavirus Disease, United States. Emerg Infect Dis 26, 1266–1273, doi:10.3201/eid2606.200516 (2020).

34 Aricescu, A. R., Lu, W. & Jones, E. Y. A time- and cost-efficient system for high-level protein production in mammalian cells. Acta Crystallogr D Biol Crystallogr 62, 1243–1250, doi:10.1107/S0907444906029799 (2006).

35 Mahmutovic, S. et al. Molecular Basis for Antibody-Mediated Neutralization of New World Hemorrhagic Fever Mammarenaviruses. Cell Host Microbe 18, 705–713, doi:10.1016/j.chom.2015.11.005 (2015).

36 Kabsch, W. Xds. Acta Crystallogr D Biol Crystallogr 66, 125–132, doi:10.1107/S0907444909047337 (2010).

37 Evans, P. R. & Murshudov, G. N. How good are my data and what is the resolution? Acta Crystallogr D Biol Crystallogr 69, 1204–1214, doi:10.1107/S0907444913000061 (2013).

38 McCoy, A. J. et al. Phaser crystallographic software. J Appl Crystallogr 40, 658–674, doi:10.1107/S0021889807021206 (2007).

39 Jones, T. A., Zou, J. Y., Cowan, S. W. & Kjeldgaard, M. Improved methods for building protein models in electron density maps and the location of errors in these models. Acta Crystallogr A 47 (Pt 2), 110–119, doi:10.1107/s0108767390010224 (1991).

40 Adams, P. D. et al. PHENIX: a comprehensive Python-based system for macromolecular structure solution. Acta Crystallogr D Biol Crystallogr 66, 213–221, doi:10.1107/S0907444909052925 (2010).

41 Bricogne, G. et al. BUSTER version 2.10.3., (Global Phasing Ltd., 2017).

42 Radoshitzky, S. R. et al. Transferrin receptor 1 is a cellular receptor for New World haemorrhagic fever arenaviruses. Nature 446, 92–96, doi:10.1038/nature05539 (2007).

43 Chen, S. et al. Genome-wide CRISPR screen in a mouse model of tumor growth and metastasis. Cell 160, 1246–1260, doi:10.1016/j.cell.2015.02.038 (2015).

44 Lan, J. et al. Structure of the SARS-CoV-2 spike receptor-binding domain bound to the ACE2 receptor. Nature 581, 215–220, doi:10.1038/s41586-020-2180-5 (2020).

45 Robert, X. & Gouet, P. Deciphering key features in protein structures with the new ENDscript server. Nucleic Acids Res 42, W320–324, doi:10.1093/nar/gku316 (2014).

46 Zhang, Q. et al. Cellular Nanosponges Inhibit SARS-CoV-2 Infectivity. Nano Lett 20, 5570–5574, doi:10.1021/acs.nanolett.0c02278 (2020).

